# Short-read whole-genome sequencing identifies the Tg(Tal1-cre/ERT)42-056Jrg transgene insertion site on mouse chromosome 4

**DOI:** 10.64898/2026.11.04.764251

**Authors:** Xiaodi Wu, Maya C. Bagish, Pablo Sánchez Vela, Alexander G. Goglia, Ross L. Levine

## Abstract

Tg(Tal1-cre/ERT)42-056Jrg, also known as HSC-Scl-CreER^T^, is a well-used transgenic allele that enables genetic manipulation in mouse hematopoietic cells. Using short-read whole-genome sequencing, we identified the transgene insertion site on chromosome 4 approximately 6 kb upstream of *Echdc2*.

**Key Point:**

- Short-read whole-genome sequencing is sufficient to identify the Tg(Tal1-cre/ERT)42-056Jrg transgene insertion site on mouse chromosome 4.

## Introduction

The transgenic allele Tg(Tal1-cre/ERT)42-056Jrg,^1^ also known as HSC-Scl-CreER^T^, has been a valuable tool for genetic manipulation in mouse hematopoietic cells. To create the transgene construct injected into zygote pronuclei, a 2.1-kb fragment encoding the tamoxifen-inducible recombinase CreER^T^ was cloned between a 2.4-kb genomic fragment (“6E5”) including *Tal1* exon 4 (and part of exon 5) and a 5.5-kb genomic fragment comprising a *Tal1* 3′-enhancer.

While the transgene was rationally designed, its random insertion into the genome has not been mapped to our knowledge. Facing unexpected difficulties in crossing this strain, we recognized that identification of the transgene insertion site would allow us both to anticipate inefficient crosses and to streamline colony management by distinguishing homozygous from hemizygous animals.

Targeted locus amplification is a proximity ligation–based method to map transgene insertion sites that reveals accompanying passenger fragments and structural variations.^2-4^ Additionally, a map-then-capture sequencing approach has been described, which employs Illumina short-read sequencing followed by Cas9-mediated target enrichment and Oxford Nanopore long-read sequencing.^5^ Given the cost of deploying those techniques, characterization of highly used transgenic mouse models remains a collective-action problem.

We set out to identify the transgene insertion site for the Tg(Tal1-cre/ERT)42-056Jrg allele in an economical manner using paired-end short-read whole-genome sequencing. Such an approach has been applied successfully to the smaller rice genome;^6,7^ for the mouse, however, it became disfavored a decade ago as throughputs typical of that era limited informative reads.^8^

## Materials and Methods

All mouse work was carried out according to institutional guidelines and under protocols approved by the Institutional Animal Care and Use Committee at Memorial Sloan Kettering Cancer Center. C57BL/6-Tg(Tal1-cre/ERT)42-056Jrg (HSC-Scl-CreER^T^) mice were crossed to B6.Cg-*Gt(ROSA)26Sor*^tm9(CAG-tdTomato)Hze^ (Ai9) tdTomato reporter mice,^9^ then backcrossed to C57BL/6J (The Jackson Laboratory) for five generations; sibling mating was carried out for 2–3 generations until all offspring carried the transgene, consistent with presumptive homozygosity.

Bone marrow from one adult female mouse was then isolated in the conventional manner. After red blood cell lysis, 5 million cells in single-cell suspension were used for genomic DNA (gDNA) extraction using the DNeasy Blood & Tissue Kit (Qiagen) per the manufacturer’s instructions. An aliquot of the gDNA sample was submitted for paired-end short-read whole-genome sequencing with 40× requested coverage, which was performed as follows:

After Quant-iT PicoGreen (Invitrogen) quantification and quality control by Agilent TapeStation, 500 ng gDNA was sheared using an LE220-plus Focused-ultrasonicator (Covaris) and a sequencing library was prepared using the KAPA HyperPrep Kit (Roche) with modifications. Briefly, the library was subjected to a 0.5× size selection using AMPure XP beads (Beckman Coulter) after post-ligation cleanup. The library was not amplified by PCR and was pooled equivolume and quantitated based on its initial sequencing performance. The sample was run on a NovaSeq X in a PE150 run using the NovaSeq X 10B Reagent Kit (Illumina). The sample was sequenced to yield 1.3 billion reads.

Using mouse genome assembly mm10 as a starting point, a custom augmented genome was generated by appending the CreER^T2^-poly(A) fragment from a published plasmid (Addgene sequence 14797) as an additional, surrogate reference contig. Reads were aligned against the custom augmented genome using bowtie2 and discordant pairs were obtained using samtools (`samtools view -bS -F 14`).

PCRs were performed using Q5 High-Fidelity 2× Master Mix (New England Biolabs), template gDNA either from the sample described above or from a control sample similarly prepared, and one of the following primer pairs: F1/R2, F1/R1, or F2/R2 (F1: 5′-AGATGTCCAGGCAATCCAGA-3′; R1: 5′-CCACAGAGAGAATCCCACCC-3′; F2: 5′-AGTGTGTGTCAGGCTGCAAA-3′; R2: 5′-CTTCGGTAGAGCAGTCAGTG-3′). Sanger sequencing (Azenta) was performed using one of the following primers: R1, R3, or F3 (R3: 5′-GTGTGAACTCAGGATGCAGGA-3′; F3: 5′-GGGTGCTGTTTGTGTGCAAT-3′).

### Data sharing

Original high-throughput sequencing data have been deposited in the NCBI Sequence Read Archive (SRA) under BioProject accession PRJNA1213845. Genotyping probes for automated real-time PCR (Transnetyx) can be shared upon request.

## Results and Discussion

We began with inspection of discordant read pairs in the aligned sequencing data wherein one read was mapped to the “6E5” fragment or the *Tal1* 3′-enhancer fragment, finding informative mates mapped to chromosome 1 (qC2) and chromosome 4 (qC7). Further inspection centered on those loci indicated the following genomic arrangement (coordinates relative to mouse genome assembly mm10): at the 5′-junction, a fragment of the *Tal1* 3′-enhancer is inserted in reverse orientation downstream of chr4:108,158,324; at the 3′-junction, the *Tal1* 3′-enhancer is inserted in forward orientation, followed by a 1.7-kb passenger fragment (chr1:61,262,976–61,264,685), all upstream of chr4:108,159,502 **(Fig. 1A)**.

**Figure 1.**
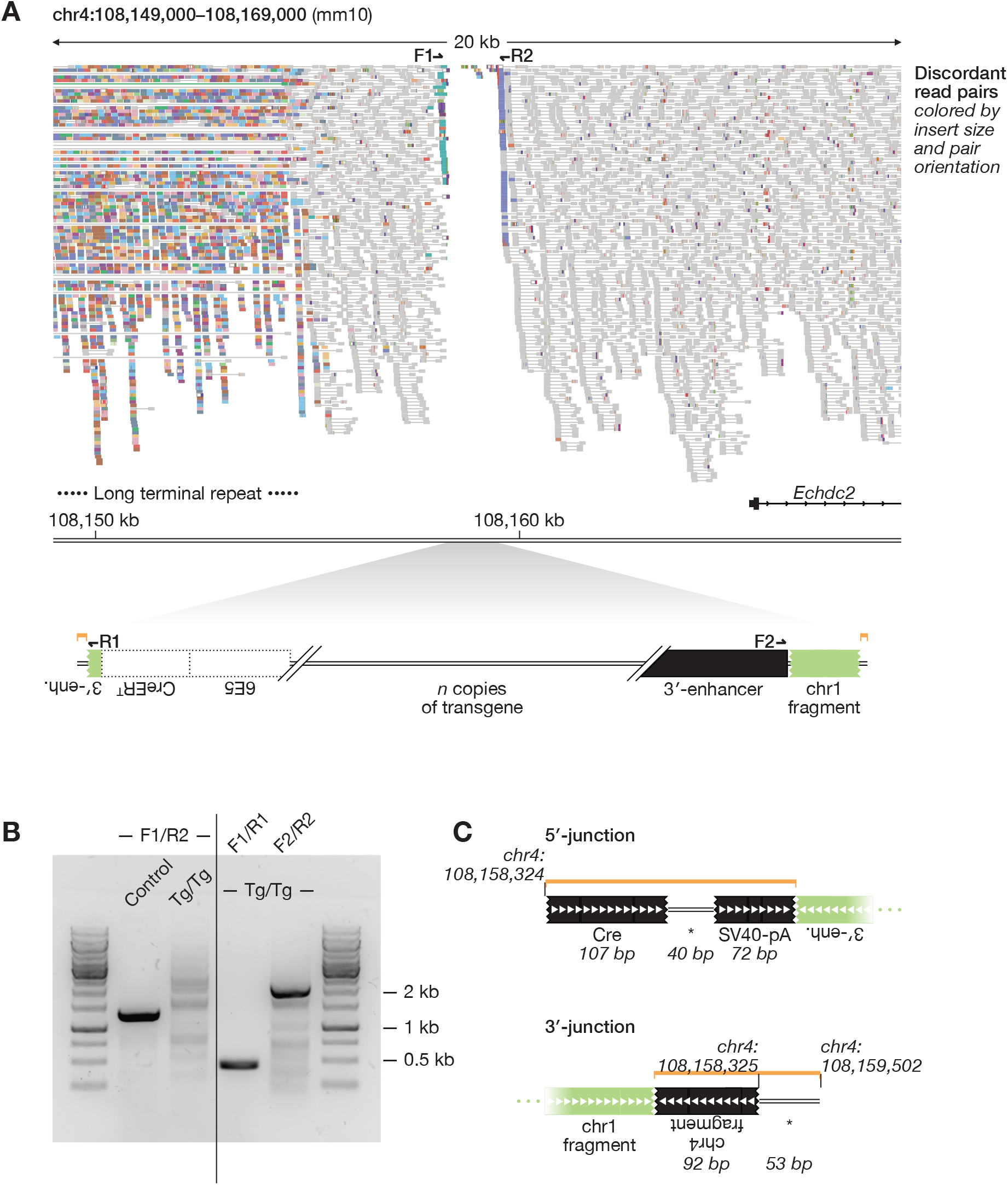
Tg(Tal1-cre/ERT)42-056Jrg is inserted on mouse chromosome 4. (A) Discordant read pairs with at least one read mapping to a 20-kb region that includes the transgene insertion site, colored by insert size and pair orientation. Teal reads left of center have mates mapped to the *Tal1* 3′-enhancer (enh., lime green at bottom left); blue-violet reads right of center have mates mapped to a region of chromosome 1 (lime green at bottom right). Other discordant read pairs with one read mapping to that region of chromosome 1 (not shown) in turn have mates mapped to the *Tal1* 3′-enhancer, consistent with a passenger fragment arranged as depicted at bottom. Harpoons represent primers referenced in (B). Jagged edges indicate premature truncation of genetic elements. Dotted outlines show the positions of genetic elements based on their arrangement in the rationally designed transgene. (B) Agarose gel electrophoresis showing PCR amplicons obtained using the indicated primer pairs and template gDNA. Control: non-transgenic mouse gDNA; Tg/Tg: putatively homozygous transgenic mouse gDNA. (C) Diagram illustrating the arrangement of small genetic fragments in regions (orange brackets) immediately adjacent to the 5′- and 3′-junctions based on Sanger sequencing results. Asterisks mark fragments of unknown provenance. SV40-pA: SV40 poly(A) signal element.

The transgene insertion site does not disrupt the coding sequence of any annotated endogenous gene, with the closest being *Echdc2* (encoding mitochondrial enoyl-CoA hydratase domain–containing protein 2) located ∼6 kb downstream. By comparison, the *Jak1* coding sequence is found ∼7 Mb upstream, while the native *Tal1* coding sequence is found ∼7 Mb downstream.

PCR using primers flanking the transgene insertion site amplified a 1.3-kb wild-type fragment (chr4:108,158,231–108,159,561) from control animal gDNA but not from putatively homozygous transgenic animal gDNA; meanwhile, PCRs using primers flanking the 5′-and 3′-junctions amplified fragments of expected size from transgenic animal gDNA **(Fig. 1B)**. Sanger sequencing of those junctions corroborated the genomic arrangement and added information about interposed small fragments **(Fig. 1C)**.

We report these findings both because they further characterize a highly used transgenic mouse model and because they redemonstrate that such work can be carried out without specialized methodologies. Using paired-end short-read whole-genome sequencing, our approach has been deliberately scoped to support practical animal husbandry needs. Insert copy number and surrounding structural variations have not been systematically analyzed: long-read sequencing with or without adaptive sampling,^10^ when affordable, will readily yield those insights one day.

## Acknowledgments

We are grateful to lab members for discussion and critical reading. XW was supported by a career development award from Blood Cancer United (formerly The Leukemia & Lymphoma Society). AGG was supported by the Memorial Sloan Kettering Louis V. Gerstner Jr. Physician Scholar Program and NCI award L30 CA294382. RLL was supported by NCI award R35 CA197594 and by a SCOR grant from Blood Cancer United. We acknowledge use of the Integrated Genomics Operation core facility, funded by the NCI Cancer Center Support Grant (P30 CA008748), Cycle for Survival, and the Marie-Josée and Henry R. Kravis Center for Molecular Oncology.

## Authorship Contributions

XW and RLL designed research, interpreted data, and wrote the manuscript. XW, MCB, and AGG collected data. PSV contributed vital reagents.

